# Astrocyte interferon-gamma signaling dampens inflammation during chronic central nervous system autoimmunity via PD-L1

**DOI:** 10.1101/2023.06.20.545679

**Authors:** Brandon C. Smith, Rachel A. Tinkey, Arshiya Mariam, Maria L. Habean, Ranjan Dutta, Jessica L. Williams

**Affiliations:** Department of Neurosciences, Lerner Research Institute, Cleveland Clinic, Cleveland, OH; Department of Biological, Geological, and Environmental Sciences, Cleveland State University, Cleveland, OH; School of Biomedical Sciences, Kent State University, Kent, OH; Department of Quantitative Health Sciences, Lerner Research Institute, Cleveland Clinic, Cleveland, OH; Department of Neurosciences, Case Western Reserve University School of Medicine, Cleveland, OH

## Abstract

Multiple sclerosis (MS) is an inflammatory and neurodegenerative disease of the central nervous system (CNS). Infiltrating inflammatory immune cells perpetuate demyelination and axonal damage in the CNS and significantly contribute to pathology and clinical deficits. While the cytokine interferon (IFN)γ is classically described as deleterious in acute CNS autoimmunity, we and others have shown astrocytic IFNγ signaling also has a neuroprotective role. Here, we performed RNA sequencing and ingenuity pathway analysis on IFNγ-treated astrocytes and found that PD-L1 was prominently expressed. Using a PD-1/PD-L1 antagonist, we determined that apoptosis was reduced in leukocytes exposed to IFNγ-treated astrocytes *in vitro*. To further elucidate the role of astrocytic IFNγ signaling on the PD-1/PD-L1 axis *in vivo*, we induced the experimental autoimmune encephalomyelitis (EAE) model of MS in *Aldh1l1-*Cre^ERT2^, *Ifngr1*^fl/fl^ mice. Mice with conditional astrocytic deletion of IFNγ receptor exhibited a reduction in PD-L1 expression which corresponded to increased infiltrating leukocytes, particularly from the myeloid lineage, and exacerbated clinical disease. PD-1 agonism reduced EAE severity and CNS-infiltrating leukocytes. Importantly, PD-1 is expressed by myeloid cells surrounding MS lesions. These data support that IFNγ signaling in astrocytes diminishes inflammation during chronic autoimmunity via upregulation of PD-L1, suggesting potential therapeutic benefit for MS patients.

## Introduction

Multiple sclerosis (MS) is a chronic inflammatory, demyelinating disease of the central nervous system (CNS) and remains one of the most common non-trauma-related disabling disorders among young adults (1–3). MS is characterized by sensory and motor deficits that result from loss of myelin and axons and is perpetuated by the infiltration of autoreactive immune cells that promote neuroinflammation (4, 5). Most patients experience a relapsing-remitting form of MS (RRMS) that is characterized by periods of neurological dysfunction followed by periods of remission. A subset of RRMS patients progress into a secondary progressive (SPMS) form in which periods of remission lessen and neurological disability is enhanced (6–8). Additionally, some patients are diagnosed with a third subtype, known as primary progressive MS (PPMS), in which there is consistent loss of neurological function without remission (7, 9, 10).

The progressive forms of MS, SPMS and PPMS, are characterized as primarily neurodegenerative (10), highlighted by the presence of chronic active lesions that are comprised of few infiltrating immune cells. Due to their immunomodulatory nature, this renders many of the current FDA-approved treatments relatively ineffective in treating SPMS and PPMS (11–14). Rather, the accumulation of activated, phagocytic myeloid cells on the lesion edge and astrocytic glial scaring are thought to promote a chronic, neuroinflammatory microenvironment that leads to slow and “smoldering” lesion expansion (15). Interestingly, these “smoldering” lesions are most prominent in chronic stages of MS and the density of microglia/macrophages increases with the chronicity of lesions and disease duration, particularly in progressive MS patients (16, 17). As a result, identifying pathways that target microglial and myeloid cells located at the chronic MS lesion border may serve as an efficacious treatment option and work to slow lesion expansion during chronic stages of disease.

The interferon (IFN)γ signaling pathway has classically been characterized as proinflammatory as it has been shown to drive inflammatory mechanisms such as T helper type 1 cell differentiation as well as activation of antigen presenting cells (18–23). However, more recent evidence suggests that IFNγ may facilitate neuroprotection during chronic stages of MS and in an animal model of MS, experimental autoimmune encephalomyelitis (EAE). In patients with progressive MS, improved symptoms correlated with higher serum levels of IFNγ (24) and intraventricular administration of IFNγ during chronic stages of EAE resulted in reduced disease severity and mortality in marmosets and rats (25, 26). Likewise, global loss of IFNγ or IFNγ receptor resulted in exacerbated EAE in several murine models (27–33). In addition, the beneficial effects of IFNγ during EAE can also be cell-type specific, as mice with constitutively deficient *Ifngr1* in astrocytes demonstrate increased disease severity, inflammation, and demyelination in chronic phases (34, 35).

Notably, programmed death 1 (PD-1) is an immune checkpoint that is highly upregulated by IFNγ to begin inflammation resolution and promote tissue repair by inducing immune cell exhaustion and death following programmed death ligand 1 (PD-L1) binding (36). Extracellular PD-1 is a single immunoglobulin-like domain, and its cytoplasmic region contains an immunoreceptor tyrosine-based inhibitory motif and an immunoreceptor tyrosine-based switch motif. Upon phosphorylation, src homology 2-domain-containing tyrosine phosphatase 1 (SHP1) and SHP2 are recruited (37). Recently, in the context of tumor immunity, PD-1 activation in macrophages was shown to inhibit phagocytosis (38), and following spinal cord injury, PD-1 promoted anti-inflammatory CNS myeloid cell polarization and improved motor function (39). Moreover, in inflammatory, active MS lesions, PD-L1 is expressed by resident glia, which controls lymphocyte numbers (40).

Despite these previous findings, how compartmentalized myeloid cells on chronic active MS lesion borders may be regulated by PD-1 activation has not yet been explored. Here, we demonstrate that IFNγ signaling in astrocytes upregulates PD-L1 expression that leads to immune cell exhaustion via PD-L1/PD-1 interactions during chronic stages of EAE. Furthermore, we demonstrate that PD-1 agonism is effective at dampening chronic disease and that PD-1 expression is relegated to myeloid cells in chronic active lesions of MS patient tissue. Taken together, these results advance our understanding of protective roles for IFNγ signaling in astrocytes and identify PD-1 agonism as a potential therapeutic modality for patients with chronic, progressive MS.

## Results

### IFNγ modulates the PD-1/PD-L1 axis in human astrocytes

Our previous work identified a beneficial role for IFNγ signaling in astrocytes during chronic CNS autoimmunity (41). Since astrocytes are present throughout all subtypes of MS lesions (42, 43) and IFNγ is present at all stages of MS (44), we further explored the transcriptional control of IFNγ signaling in astrocytes. RNA-sequencing of IFNγ-treated primary human spinal cord astrocytes revealed a significant number of differentially expressed genes compared to untreated astrocytes via ingenuity pathway analysis (IPA) (Figure 1A). IPA identified significant changes in the antigen presentation pathway, which we explored previously (41), and in several others including the Th1, Th1 and Th2, PD-1/PD-L1 cancer immunotherapy, and tumor microenvironment pathways, all of which included genes relating to the PD-1/PD-L1 signaling axis (Figure 1A). Notably, in the PD-1/PD-L1 cancer immunotherapy pathway, *CD274*, the gene that transcribes PD-L1, *PDCD1LG2*, the gene that transcribes PD-L2, and *PTPN11*, the gene that encodes SHP2 are all predicted to be activated by IFNγ in astrocytes (Figure 1B). Further RNA-sequencing analysis of astrocyte *CD274*, *PDCD1LG2*, and *PTPN11* revealed that IFNγ stimulation for 24 h increased expression levels of *CD274* and *PDCD1LG2*, but did not change *PTPN11* expression (Figure 1C). These data suggest that IFNγ signaling in human spinal cord astrocytes upregulates several potential pathways that may work to limit neuroinflammation, including genes in the PD-1/PD-L1 axis, but that astrocytes themselves likely do not undergo PD-1-mediated cell death.

**Figure 1.**
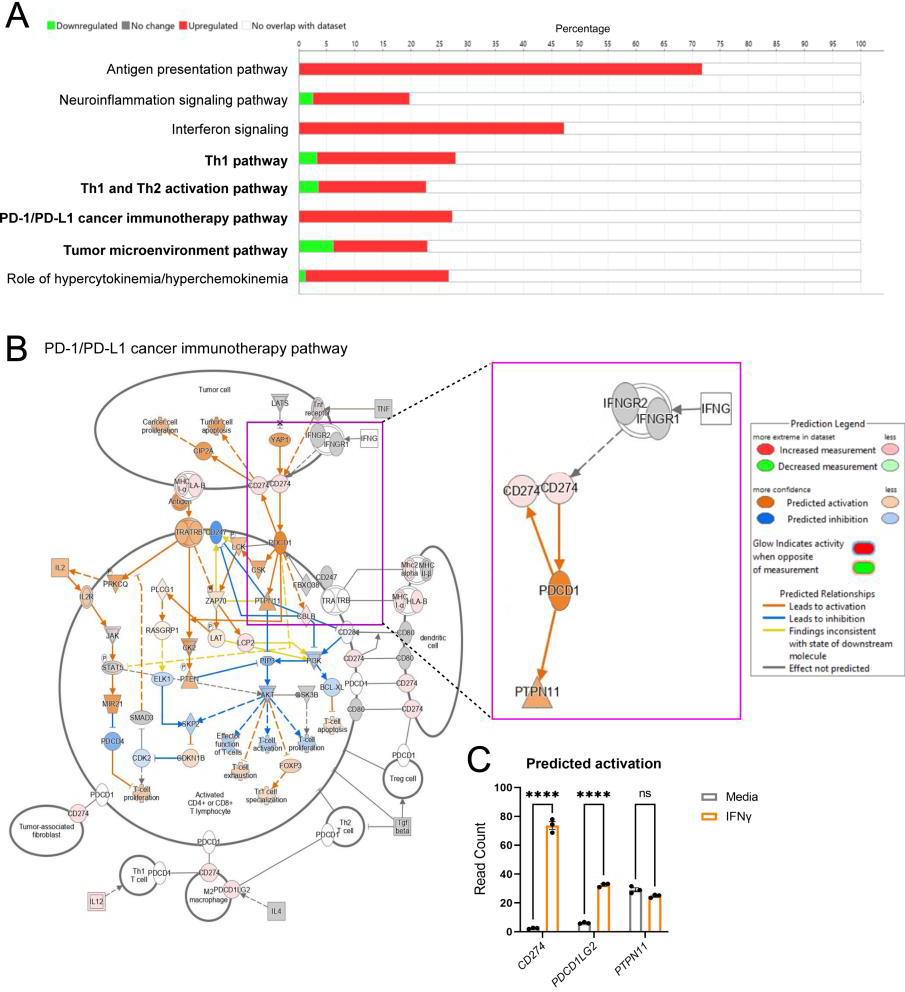
IFNγ signaling induces the PD-1/PD-L1 axis in astrocytes. Modulated Ingenuity Pathway Analysis (**A**) pathways and (**B**) signaling networks 24-h stimulation of primary human astrocytes with 10 ng/ml IFNγ and RNA sequencing. (**C**) A two-way ANOVA with Tukey’s multiple comparison test was performed on read counts of differentially expressed genes relevant to the PD-1/PD-L1 axis. Data represent the mean ± SEM from 3 independent samples. ****P < 0.0001.

### IFNγ-mediated astrocyte PD-L1 expression induces leukocyte apoptosis

We further validated the transcriptional regulation of PD-1 and PD-L1 in astrocytes by IFNγ using quantitative real time PCR. Transcript levels of PD-L1 were significantly increased in both human and murine astrocytes treated with IFNγ, while PD-1 expression levels remained unchanged relative to media controls (Figure 2A, B). Similarly, PD-L1 protein expression was enhanced in human and murine primary spinal cord astrocytes following treatment with IFNγ (Figure 2C-F). These data suggest that IFNγ signaling induces astrocyte immune checkpoint expression, while astrocytes themselves do not undergo apoptosis due to the lack of PD-1 expression. The role of PD-L1 as an immune checkpoint is well established (45, 46); however, whether astrocytic PD-L1 directly induces apoptosis in leukocytes has not been shown. To determine the functional capacity of PD-L1 on astrocytes to act as an immune checkpoint, we cocultured primary murine astrocytes and lymph node cells (LNCs) for 48 h in the presence or absence of IFNγ, a PD-1/PD-L1 inhibitor, a PD-1 agonist, IFNγ + the PD-1/PD-L1 inhibitor, or IFNγ + PD-1 agonist. Following coculture, LNCs were mechanically removed from astrocytes, counted, and LNCs were subjected to a caspase 3/7 apoptosis activity assay (Figure 2G). Relative to media controls, the LNCs that were cocultured with astrocytes had an increase in apoptosis when treated with IFNγ and/or the PD-1 agonist. Importantly, this increase was diminished when cells were treated with the PD-1/PD-L1 inhibitor (Figure 2H). These data suggest that IFNγ-mediated PD-L1 expression on astrocytes is functional and likely acts as a traditional immune checkpoint. To determine if astrocytes were necessary for the induction of apoptosis, LNCs were cultured and treated alone, in the absence of astrocytes. As anticipated, only LNCs treated with the PD-1 agonist exhibited evidence of Caspase 3/7 activity (**Supplemental Figure 1A**). To determine how IFNγ, the PD-1 agonist, or PD-1 antagonist affected astrocyte caspase activity, astrocytes were cultured with and without LNCs and treated. Following the removal of LNCs or when astrocytes were cultured alone, there was no change in astrocyte apoptotic activity observed (**Supplemental Figure 1B, C**). As an additional blank control to confirm the lack of nonspecific Caspase 3/7 activity, LNCs were cultured, treated, and removed (**Supplemental Figure 1D**). Finally, qPCR analysis of LNCs treated with and without IFNγ resulted in no change in transcript levels of *Cd274* or *Pdcd1* (**Supplemental Figure 1E**). Taken together, these data suggest that astrocytes do not self-regulate via PD-1/PD-L1 signaling, but rather the expression of PD-L1 on astrocytes influences the apoptotic activity of neighboring LNCs.

**Figure 2.**
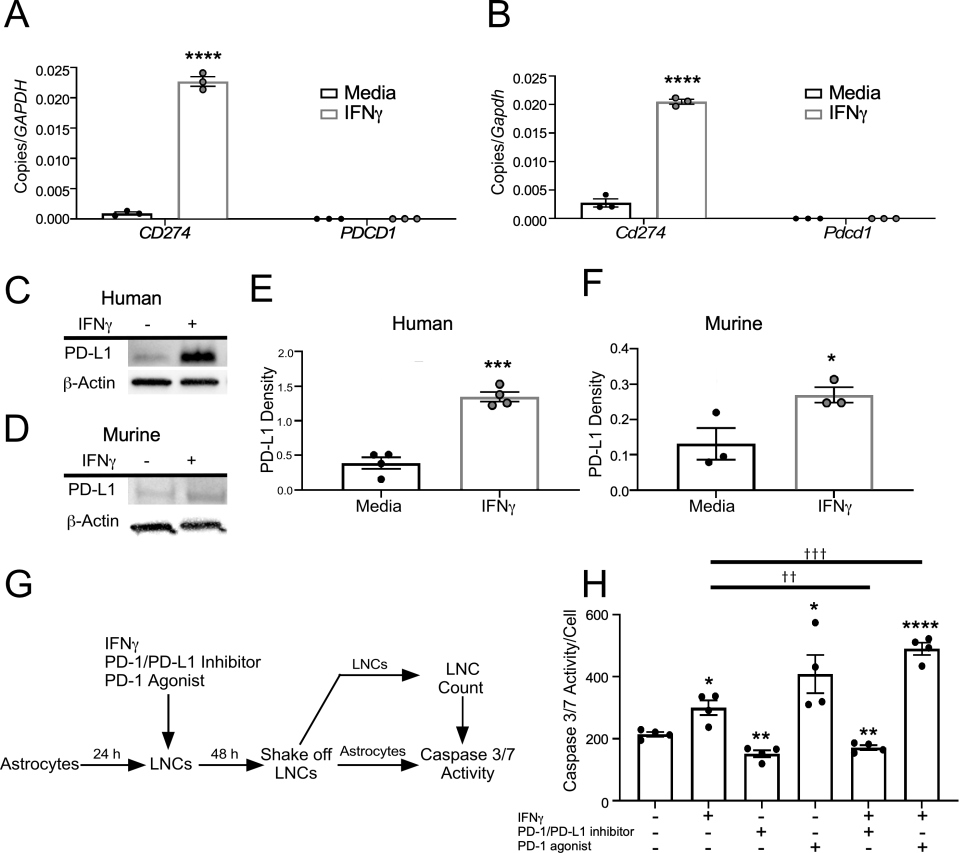
IFNγ-regulated expression and function of PD-L1 in astrocytes. (**A**) Primary human spinal cord astrocytes and (**B**) primary murine spinal cord astrocytes were stimulated with and without 10 ng/ml IFNγ for 24 h and RNA was collected and analyzed for transcript levels of *CD274* and *PDCD1* by qRT PCR. (**C**) Primary human spinal cord and (**D**) primary murine spinal cord astrocytes were stimulated with or without 10 ng/ml IFNγ for 24 h and protein lysates were assessed for expression of PD-L1 via Western blot. (**E, F**) PD-L1 levels were quantified and normalized to β-actin expression. (**G**) Schematic representation of the experimental design for data presented in panel H. (**H**) Caspase 3/7 activity was quantified and normalized to lymph node cell number following coculture with astrocytes treated with 10 ng/ml IFNγ. Data are representative of 2 independent experiments with 3-4 technical replicates each. All data represent the mean ± SEM. *P < 0.05, **P < 0.01, ***P < 0.001, ****P < 0.0001 compared to media-treated samples and ^††^P < 0.01, ^†††^P < 0.001 between treatments by two-way ANOVA.

### Astrocytic IFNγ signaling induces PD-L1 expression in astrocytes and modulates lesion composition during chronic EAE

We and others have shown that disrupted IFNγ signaling in astrocytes exacerbates chronic EAE (35, 41). Here, we extend those findings and demonstrate that astrocyte PD-L1 expression is downstream of IFNγ signaling in astrocytes and has a role in controlling CNS-infiltrating leukocyte populations during the later stages of EAE. To examine how IFNγ signaling in astrocytes impacted PD-L1 expression during EAE, we immunized *Ifngr1*^fl/fl^ *Aldh1l1-*Cre^ERT2+^ mice and littermate controls and starting one day following peak disease, mice were injected with tamoxifen to conditionally delete the IFNγ receptor from astrocytes (**Supplemental Figure 2**). Following tamoxifen administration, *Ifngr1*^fl/fl^ *Aldh1l1-*Cre^ERT2+^ mice exhibited exacerbated EAE, a larger lesion burden, and greater myelin loss compared to littermate controls (Figure 3A, **Supplemental Figure 3**). Importantly, there was a significant decrease in both total PD-L1 expression and PD-L1 colocalized with GFAP^+^ astrocytes within the lesions of *Ifngr1*^fl/fl^ *Aldh1l1-*Cre^ERT2+^ mice compared to littermate controls (Figure 3B-E). Consistent with this, IHC analysis of ventral spinal cord white matter tracts revealed that compared to controls, *Ifngr1*^fl/fl^ *Aldh1l1-*Cre^ERT2+^ mice had increased PD-1 expressing cells within lesions (Figure 3F-L). We found that mice lacking intact astrocytic IFNγ signaling had increased T cell infiltration (Figure 3M); and, in examining CD45^+^ and Iba1^+^ cell populations, we found that lesions in *Ifngr1*^fl/fl^ *Aldh1l1-*Cre^ERT2+^ mice contained significantly enhanced populations of Iba1^+^CD45^+^ cells compared to controls (Figure 3N). To ensure that manipulation of IFNγ signaling on astrocytes was not altering PD-1 expression on targeted cells, we determined PD-1 colocalization with CD3^+^, Iba1^+^, and GFAP^+^ cells and found no observable difference (Figure 3O). Together, these data suggest that during chronic EAE, IFNγ signaling enhances PD-L1 expression on spinal cord astrocytes which leads to a decrease in infiltrating leukocytes, particularly activated cells of the myeloid lineage.

**Figure 3.**
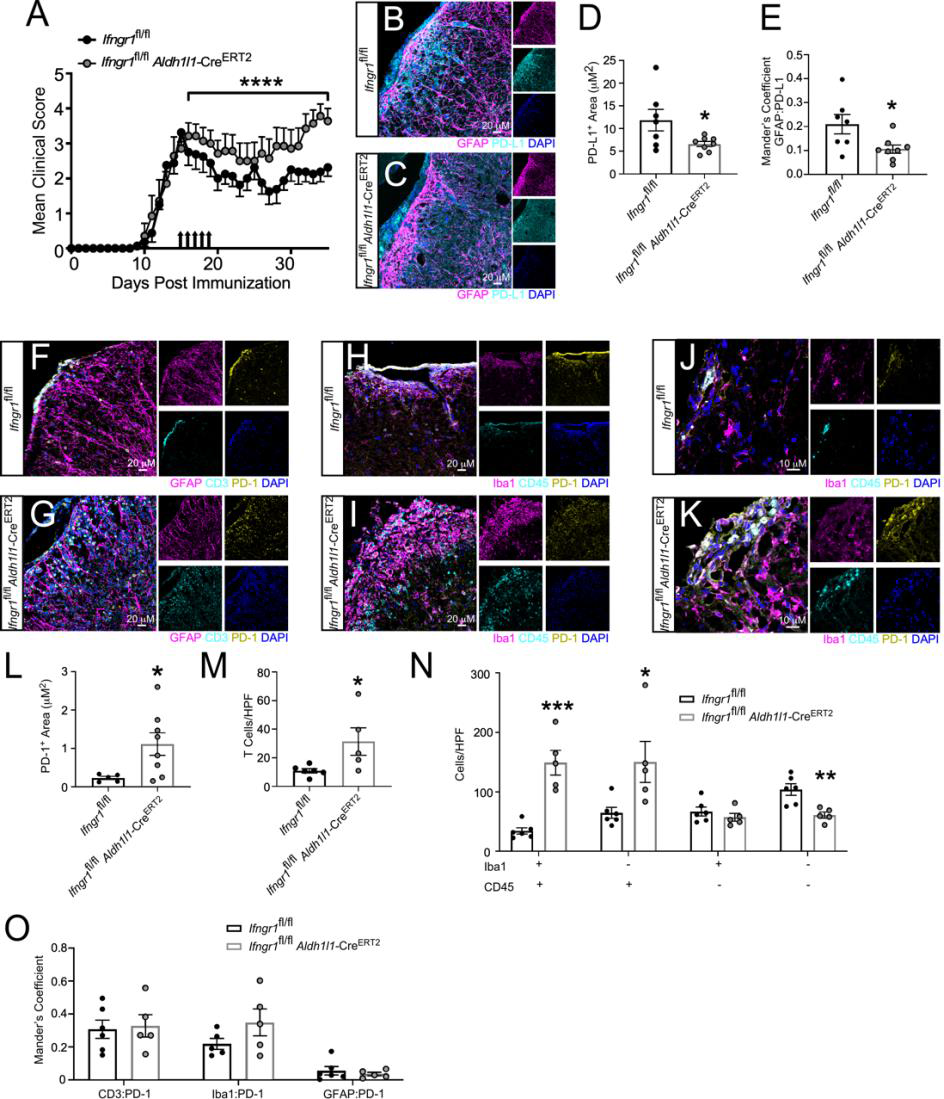
Astrocyte IFNγ signaling is protective and upregulates PD-L1 during chronic EAE. (**A**) EAE was induced in *Ifngr1*^fl/fl^ *Aldh1l1*-Cre^ERT2+^ mice (*n* = 7) and *Ifngr1*^fl/fl^ littermate controls (*n* = 8) and EAE clinical course was blindly monitored. On day 16 ± 1 mice were injected i.p. with tamoxifen for 5 consecutive days to induce recombination (black arrows). Graph is representative of two combined independent experiments. 35 days post-immunization, mice were perfused and the CNS was removed and cryopreserved for IHC analysis. Ventral white matter tracts of the lumbar spinal cord were imaged using confocal microscopy. (**B**) *Ifngr1*^fl/fl^ and (**C**) *Ifngr1*^fl/fl^ *Aldh1l1*-Cre^ERT2+^ tissue sections were labeled for GFAP, PD-L1, and nuclei were counterstained with DAPI. (**D**) Total PD-L1 area and (**E**) PD-L1 colocalized with GFAP were analyzed using ImageJ. (**F, G**) Tissue sections were labeled for GFAP, CD3, and PD-1 and (**H-K**) for Iba1, CD45, PD-1. Nuclei were counterstained with DAPI and imaged at (**H, I**) 20x and (**J, K**) 63x magnification. (**L**) Total PD-1 area, (**M**) CD3^+^ cells and (**N**) Iba1^+^ and CD45^+^ cells per high powered field were quantified. (**O**) Colocalization of PD-1 was assessed for CD3^+^, Iba1^+^, and GFAP^+^ cells using ImageJ. Data in panel A represent the mean ± SEM and were analyzed using a Mann–Whitney *U* test for nonparametric data. Data represent the mean ± SEM and were analyzed using a two-tailed Student’s *t* test (D, E, L, M) or two-way ANOVA (N, O). Data are combined from two independent experiments. *P < 0.05, **P < 0.01, ***P < 0.001.

### PD-1 agonism reduces chronic EAE severity

While CNS infiltration is not as prominent during chronic EAE as compared to acute stages, there is still an appreciable level of neuroinflammation (47–49). The role of PD-1 agonism in dampening these remaining CNS infiltrating populations during chronic EAE has yet to be explored. To address this, we induced EAE in WT C57Bl/6 mice and two days after peak disease, we began administering PD-1 agonist or a vehicle control for 5 consecutive days. Following treatment, EAE progression ceased in mice receiving PD-1 agonist compared to vehicle-treated mice (Figure 4A). We determined that PD-1 agonism had no impact on astrocyte reactivity by IHC labeling; however, we did observe an increase in myelin basic protein (MBP)^+^ area and a corresponding decrease in lesion size in agonist-treated mice compared to vehicle controls (Figure 4B, C, H-J). To determine if canonical exhaustion pathways were activated following PD-1 agonism, we labeled the spinal cords of treated mice for phosphorylated-SHP2 (pSHP2) (50). Indeed, IHC labeling of PD-1 agonist-treated mice revealed an overall increase in pSHP2^+^ area, despite a decrease in CD11b^+^ cells (Figure 4D-G, K, L). Finally, using higher magnification for analysis of colocalization, we found a significant increase in pSHP2 colocalized with CD11b following PD-1 agonism compared to vehicle treatment (Figure 4F, G, M). These data suggest that PD-1 agonism has a potential role in limiting myeloid cell-mediated CNS inflammation during chronic autoimmunity.

**Figure 4.**
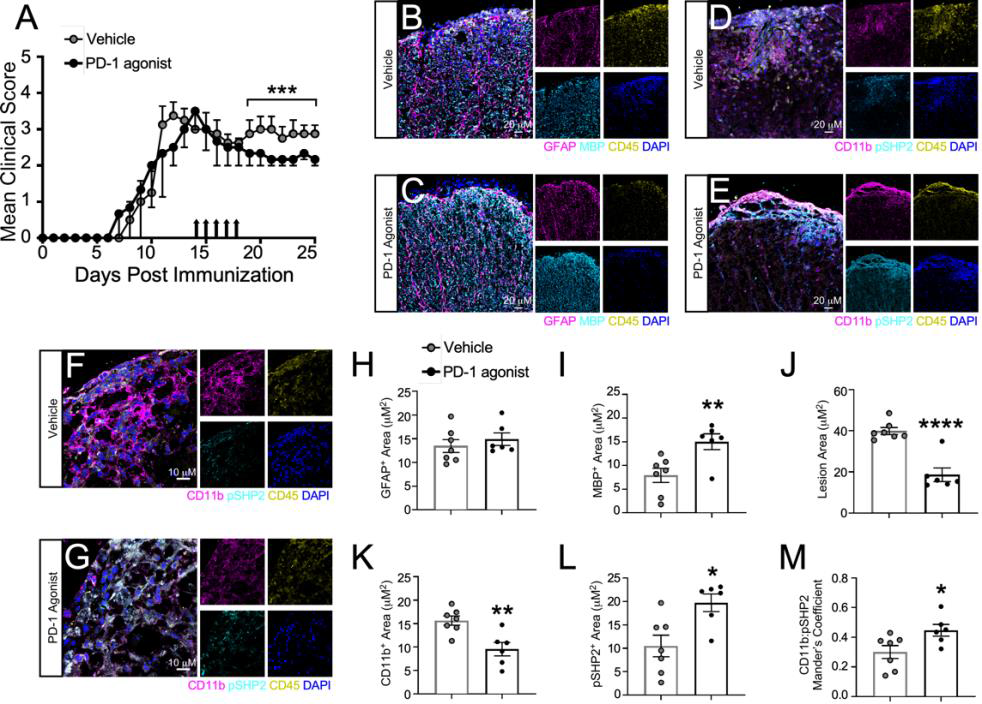
PD-1 agonism prevents the progression of EAE. (**A**) EAE was induced in WT C57Bl/6J mice. Clinical course was blindly monitored. One day after peak disease, a PD-1 agonist (*n* = 6) or vehicle control (*n* = 6) treatment was randomly assigned and injected i.p. for 5 consecutive days. Data are representative of two independent experiments. (**B-M**) 25 days post-immunization, mice were sacrificed and the CNS was collected and cryopreserved for IHC analysis. Ventral white matter tracts of the lumbar spinal cord were imaged using confocal microscopy. Spinal cord tissue from (**B**) vehicle- and (**C**) PD-1 agonist-treated mice were labeled for GFAP, MBP, CD45, and nuclei were counterstained with DAPI. Tissues from (**D, F**) vehicle- and (**E, G**) PD-1 agonist-treated mice were also labeled for CD11b, pSHP2, CD45, and nuclei were counterstained with DAPI. Total (**H**) GFAP, (**I**) MBP, (**J**) lesion area, (**K**) CD11b, and (**L**) pSHP2 were quantified using ImageJ. (**M**) Colocalization of CD11b with pSHP2 was also assessed. Data represent the mean ± SEM combined from 2 independent experiments and were analyzed using the (**A**) Mann–Whitney *U* test for nonparametric data or by (**H-M**) two-tailed Student’s *t* test. *P < 0.05, **P < 0.01, ****P < 0.0001.

### PD-1 is expressed on the rim of chronic active MS lesions

Chronic active lesions are common in chronic, progressive stages of MS (51, 52). These chronic active lesions are ringed by a dense population of microglia (52, 53). The presence of PD-1 in chronic active lesions has yet to be explored. Using post-mortem MS tissue (Table 1), we identified chronic active lesions and normal-appearing white matter (NAWM) based on the presence (or absence) of MBP and the patterning of Iba1^+^ cells (Figure 5A, B). We then examined NAWM and the lesion rim of chronic active lesions using immunofluorescence for PD-1 expression (Figure 5C-G) and found that PD-1 was significantly increased in the lesion rim compared to NAWM (Figure 5H). Colocalization analysis revealed that PD-1 expression was specific to Iba1^+^ cells within the lesion rim and was largely excluded from both NAWM- and lesion-associated GFAP^+^ astrocytes (Figure 5I). These data suggest that myeloid cells on the lesion rim of chronic active MS lesions, which are thought to contribute to lesion expansion, are potentially poised for PD-1 agonist targeting to locally dampen neuroinflammation and prevent disease exacerbation.

**Table 1.**
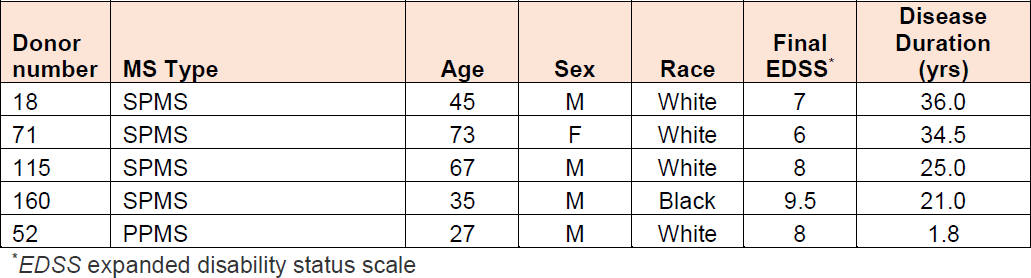
Patient Characteristics.

| **Donor number** | **MS Type** | **Age** | **Sex** | **Race** | **Final EDSS\*** | **Disease Duration (yrs)** |
| --- | --- | --- | --- | --- | --- | --- |
| 18 | SPMS | 45 | M | White | 7 | 36.0 |
| 71 | SPMS | 73 | F | White | 6 | 34.5 |
| 115 | SPMS | 67 | M | White | 8 | 25.0 |
| 160 | SPMS | 35 | M | Black | 9.5 | 21.0 |
| 52 | PPMS | 27 | M | White | 8 | 1.8 |

\*EDSS expanded disability status scale

**Figure 5.**
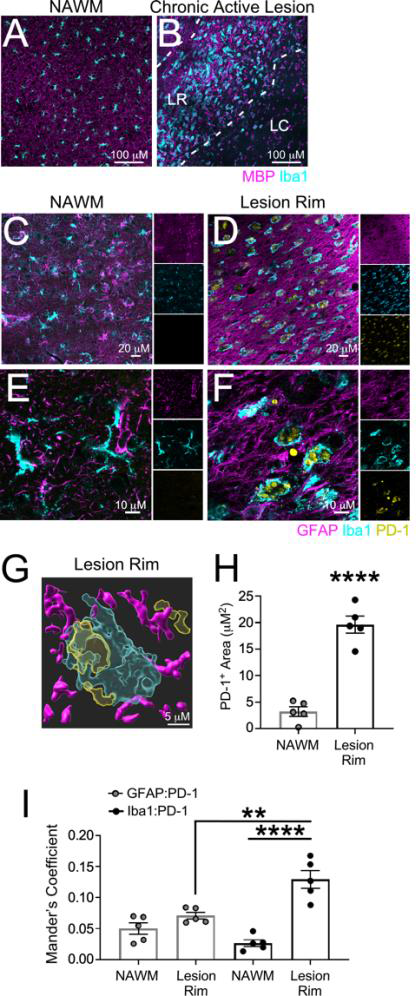
Myeloid cells at the chronic active lesion rim express PD-1. Using human post-mortem MS tissue (*n* = 5), (**A**) NAWM and (**B**) chronic active lesions were cryopreserved, sectioned, and labeled with MBP and Iba1 and imaged using confocal microscopy. Once NAWM, chronic active lesion rims (LR), and the lesion core (LC) were identified, (**C, E**) NAWM and (**D, F**) lesion rims were labeled for GFAP, Iba1, and PD-1 at (**C, D**) 20x and (**E, F**) 63x magnification using confocal microscopy. (**G**) IMARIS rendering of GFAP, Iba1, and PD-1 within the chronic active lesion rim was generated to identify PD-1 surface expression. (**H**) Total PD-1 area was quantified and (**I**) the colocalization of GFAP and Iba1 with PD-1 in both NAWM and the chronic active lesion rim was assessed. Data represent the mean ± SEM and were analyzed using a (**H**) two-tailed Student’s *t* test or (**I**) one-way ANOVA. **P < 0.01, ****P < 0.0001.

## Discussion

Increasing evidence points to a protective role for IFNγ in chronic stages of both MS and EAE (41, 44, 54). In this study, we contribute to these findings and demonstrate that IFNγ signaling in astrocytes upregulates PD-L1, a key immune checkpoint, implicating PD-L1 as contributor in IFNγ-mediated dampening of chronic neuroinflammation. Our study revealed that IFNγ-mediated upregulation of PD-L1 on astrocytes corresponded to a reduction in infiltrating leukocytes, particularly cells within the myeloid compartment, and that loss of IFNγ signaling during later stages of EAE exacerbated clinical severity. Furthermore, PD-1 agonism *in vivo* led to ameliorated EAE, with reduced CNS immune cell infiltration. This agonism resulted in increased classical downstream PD-1 exhaustion markers in the myeloid compartment. Additionally, postmortem MS tissue analysis revealed PD-1 to be concentrated on myeloid cell “rim of microglia” surrounding chronic active MS lesions.

These findings strengthen the importance of IFNγ signaling as a protective factor during chronic MS and highlight the need for unique treatment strategies for chronic MS patients. Previous studies have shown that constitutive loss of PD-1/PD-L1 during EAE leads to disease exacerbation (55). However, PD-L1 is expressed on a wide array of cell types implicated in both MS and EAE, including, but not limited to, T cells, microglia, and neurons (45, 46, 56). Our work implicates astrocytes as key drivers of the PD-1/PD-L1 axis. Furthermore, while previous studies have examined the role of PD-L1 in acute autoimmunity, our study suggests that activation of the PD-1 pathway halts the progression of chronic autoimmunity. The cellular and inflammatory makeup differ between acute and chronic MS and EAE (57–60). As such, our findings uncovering the cell type and potential kinetic activity of PD-L1 could greatly increase our understanding of mechanisms that contribute to continued and progressive disease activity seen in chronic MS patients.

Previous studies have shown the importance of PD-L1 regulation of T cells (38, 40, 56, 61). While our study recapitulates these findings to an extent, we show here that PD-L1 may also alter the myeloid compartment during chronic autoimmunity. There is a wide range of macrophage activation states. Highly activated and inflammatory macrophages tend to be pathogenic, while homeostatic macrophages are more protective (62). Macrophages are known to express PD-1 at many stages of activation (38, 63, 64), which may explain the reduction in CNS myeloid cells in *Ifngr1*^fl/fl^ *Aldh1l1-*Cre^ERT2+^ mice compared to littermate controls during the later stages of EAE and the increase in myeloid-associated pSHP2 following PD-1 agonism. Since phosphorylation of SHP2 is classically thought of as an initiating step of the PD-1/PD-L1 cell exhaustion pathway (50), the increase in pSHP2 that we find in the CNS myeloid compartment after treatment with a PD-1 agonist suggests that the remaining cells potentially have reduced activation. The post-acute timing of PD-1 agonism during the course of EAE may explain the exaggerated reduction we see in the myeloid compartment relative to the lymphoid compartment, as lymphocytes are already drastically reduced in the chronic stages of EAE relative to acute time points (47). These data suggest that myeloid cells significantly contribute to the pathophysiology of chronic EAE and that PD-1 agonism may have a role in halting chronic neuroinflammation via the reduction of the number and activation of cells in the myeloid compartment.

Activated microglia can upregulate both PD-1 and PD-L1 in the context of neurodegeneration (65). Disease-associated microglia (DAM) have a transcriptional profile that is highly inflammatory (66). These DAMs are found in EAE lesions (67) and in chronic active lesions within MS patients (67, 68). These DAMs are also a potential target for PD-1 agonism to reduce their inflammatory profile. Similar to our findings, a recent study reported that microglial PD-1 stimulation leads to reduced inflammation in Alzheimer’s disease (65). Of note, Kummer and colleagues postulated that PD-L1 shedding contributed to the efficaciousness of targeting the PD-1/PD-L1 axis (65). While we did not examine PD-L1 shedding here, it is likely that shedding would result in increased PD-1 signaling, further dampening autoimmune neuroinflammation. Since myeloid cells seem to be a prominent contributor to pathology and expressors of PD-1 during chronic MS, PD-1 agonism proves to be a viable option for SPMS patients, a population of patients with highly limited treatment availability. Further, since PD-1 agonism is a strategy currently in Phase I clinical trials for rheumatoid arthritis, alopecia, and transplant rejection (69), our study provides critical evidence in moving towards treatment availability to these MS patients.

## Materials and Methods

### Pathway analysis

Total RNA was collected from human spinal cord astrocytes (ScienCell) treated with and without recombinant human IFNγ (Peprotech) for 24 h using an RNeasy Kit (QIAGEN) according to the manufacturer’s instructions. RNA was then sequenced at the Cleveland Clinic Lerner Research Institute’s Genomics Core. The dataset was then analyzed using IPA software (Ingenuity System Inc. USA) to examine the canonical pathways upregulated by IFNγ compared to media-treated controls.

### Astrocytes

Primary adult human spinal cord astrocytes were obtained from ScienCell Laboratories and grown according to provided protocols in complete ScienCell Astrocyte Medium. Briefly, primary human astrocytes were isolated from the spinal cord and at P0 were tested for morphology by phase contrast and relief contrast microscopy and GFAP positivity by immunofluorescence. Cell number, viability (≥ 70%), and proliferative potential (≥ 15 pd) were also assessed, and negative screening for potential biological contaminants was confirmed prior to cryopreservation and receipt of frozen cells at P1. Primary murine spinal cord astrocytes were collected as previously described (70) from C57Bl/6J P2-4 pups.

### qRT-PCR analysis

Total RNA was collected from treated human or murine primary spinal cord astrocytes using an RNeasy Kit (QIAGEN) according to the manufacturer’s instructions. Reverse transcription and SYBR Green qRT-PCR were performed as previously described using primers specific for human *CD274* (forward: CCA AGG CGC AGA TCA AAG AGA, reverse: AGG ACC CAG ACT AGC AGC A), *PDCD1* (forward: CCA GGA TGG TTC TTA GAC TCC C, reverse: TTT AGC ACG AAG CTC either rapidly frozen for biochemical TCC GAT), and murine *Cd274* (forward: GCT CCA AAG GAC TTG TAC GTG, reverse: TGA TCT GAA GGG CAG CAT TTC), and *Pdcd1* (forward: ACC CTG GTC ATT CAC TTG GG, reverse: CAT TTG CTC CCT CTG ACA CTG) (41). Transcript levels were normalized to copies of human *GAPDH* (forward: GAA GGT GAA GGT CGG AGT C, reverse: GAA GAT GGT GAT GGG ATT TC) or murine *Gapdh* (forward: GGC AAA TTC AAC GGC ACA GT, reverse: AGA TGG TGA TGG GCT TCC C), respectively.

### Western blotting

Protein lysates were collected from primary human and murine spinal cord astrocytes in radioimmunoprecipitation assay (RIPA) buffer (Sigma-Aldrich) supplemented with a protease and phosphatase-3 inhibitor cocktail (Sigma-Aldrich), then 20 μg of protein was resolved on a 4-12% Tris gel and transferred to a polyvinylidene difluoride (PVDF) membrane using the Trans-Blot Turbo system (Bio-Rad) according to standard protocols. Membranes were incubated overnight at 4°C in Tris-buffered saline, 0.1% Tween® 20 (TBST), and 5% powdered milk. Membranes were blotted with anti-human PD-L1 (Invitrogen; 14-9969-82), anti-mouse PD-L1 (BioLegend; 135202), and anti-β-actin (ThermoFisher Scientific; MA5-15739) antibodies, washed with TBST 3 times, and then incubated with HRP-conjugated secondary antibodies (ThermoFisher Scientific) for 1 h at room temperature. Membranes were washed with TBST 3 times and imaged using the ChemiDoc MP imaging system (Bio-Rad) after activation with ECL substrate solution.

### Apoptosis assay

Primary murine astrocytes were cultured to 60-70% confluency in a 96-well plate for 24 h. Astrocytes were then cultured with and without 14 x 10^3^ LNCs/well and treated with 10 ng/mL recombinant murine IFNγ (Peprotech), 100 nM PD-1/PD-L1 inhibitor (Thomas Scientific; C790F18), and/or 1.0 μg/mL PD-1 agonist (BioLegend; 758204) for 48 h. LNCs were unadhered using an orbital shaker at 180 rpm for 2 h. Caspase activity was quantified using the Caspase-Glo 3/7 Assay Kit (Promega).

### EAE induction

Mice of mixed sex were induced for EAE at 8–10 weeks of age. *Aldh1l1-*Cre^ERT2^, *Ifngr1*^fl/fl^, and C57Bl/6J wild-type mice were obtained commercially from The Jackson Laboratory and housed under specific pathogen-free conditions. Mice were crossed according to standard breeding schemes to generate *Ifngr1*^fl/fl^ *Aldh1l1-*Cre^ERT2^ and littermate controls. On day 0, mice were immunized s.c. with 100 μg MOG_35-55_ emulsified in complete Freund’s adjuvant containing 400 mg heat killed Mycobacterium tuberculosis H37Ra using a standard emulsion (Hooke Laboratories). Pertussis toxin (100 ng) (Hooke Laboratories) was injected i.p. on the day of immunization and 2 days later. Mice were monitored daily for clinical signs of disease as follows: 0, no observable signs; 1, limp tail; 2, limp tail and ataxia; 2.5, limp tail and knuckling of at least one limb; 3, paralysis of one limb; 3.5; partial paralysis of one limb and complete paralysis of the other; 4, complete hindlimb paralysis; 4.5, moribund; 5, death. Tamoxifen (75 mg/kg) (Sigma) was dissolved in corn oil (Sigma) and injected i.p. for 5 consecutive days to induce recombination starting at one day post-peak EAE. Animals that did not develop clinical signs of EAE were excluded from the study.

### Immunofluorescent labeling

Mice were intracardially perfused with PBS followed by 4% paraformaldehyde (PFA) and CNS tissue was removed and fixed in 4% PFA at 4°C for 24 h. Tissue was then cryopreserved in 30% sucrose and frozen in O.C.T. Compound (Fisher HealthCare). Frozen, transverse sections (10 μm) were slide-mounted and stored at −80°C. Tissue sections were blocked with 10% goat serum and 0.1% Triton X-100 (Southern Biotech) for 1 h at room temperature and then incubated with anti-MBP (Abcam; ab7349), anti-GFAP (Invitrogen; 13-0300), -Iba1 (Wako Chemicals; 019-19741), -PD-1 (BioLegend; 135202), -CD3e (ThermoFisher; 14-0031-82), -PD-L1 (BioLegend; 124301), -CD45 PerCP-Cy5.5 (BioLegend; 103132), and/or -pSHP2 (Abcam; ab62322) primary antibodies overnight at 4°C. Secondary antibodies conjugated to Alexa Fluor 488, Alexa Fluor 555, or Alexa Fluor 647 (ThermoFisher Scientific) were applied for 1 h at room temperature as appropriate. Nuclei were counterstained with DAPI (ThermoFisher Scientific) diluted in PBS. Sections were analyzed using the 10x, 20x, or 63x objectives of a confocal microscope LSM 800 (Carl Zeiss). Images shown are representative of 3–7 images taken across two tissue sections at least 100 μm apart per individual mouse. The mean positive area, intensity, and Mander’s coefficient of colocalization were determined by setting thresholds using appropriate controls and quantified using ImageJ software (NIH).

Human periventricular white matter from MS patients (Table 1) was collected according to the established rapid autopsy protocol approved by the Cleveland Clinic Institutional Review Board. Patient tissue was removed, fixed in 4% paraformaldehyde, and sectioned for IHC analysis. Demyelinated lesions were identified and characterized by immunostaining free floating sections with MBP and Iba1 as described previously (71). Subsequent sections were used for the identification of astrocytes, myeloid cells, and PD-1. Antigen retrieval was performed by boiling tissue briefly in 10 μM citrate buffer. Sections were blocked with 5% goat serum and 0.03% Triton X-100 (Sigma-Aldrich) for 1 h at room temperature and then exposed to antibodies specific for human PD-1 (ThermoFisher Scientific; 14-9969-82), Iba1 (Wako Chemicals; 019-19741), and GFAP (Invitrogen; 13-0300) for 4-5 days at 4°C. Sections were then washed with PBS-Triton-X-100, and secondary antibodies conjugated to Alexa Fluor 488, 555, and 647 (ThermoFisher Scientific) were applied for 1 h at room temperature. Sections were then treated with 0.3% Sudan black in 70% ethanol for 3 min, imaged using the 10x, 20x, and 63x objectives of a confocal microscope LSM 800 (Carl Zeiss) and analyzed using ImageJ (NIH).

### Statistics

EAE data were analyzed using the nonparametric Mann-Whitney *U* test. Other normally distributed data were analyzed with parametric tests (2-tailed Student’s *t* test or two-way analysis of variance (ANOVA) with correction for multiple comparisons where appropriate. All statistical analyses were performed using GraphPad Prism Version 7 software (GraphPad). A *P* value of less than 0.05 was considered statistically significant. Data points in graphs represent individuals.

### Study approval

All murine procedures were approved by the Institutional Animal Care and Use Committee at the Lerner Research Institute, Cleveland Clinic Foundation (Cleveland, OH) using protocol numbers 1862 and 1871. All mice used were on a C57BL/6J background, were procured from The Jackson Laboratories and maintained on a 12-h light/dark cycle and had *ad libitum* access to food and water. Mice (housed 2-5/cage) did not have any prior history of drug administration, surgery or behavioral testing.

All human tissue used was collected as part of the tissue procurement program approved by the Cleveland Clinic Institutional Review Board. MS patient brain tissue was collected according to a rapid autopsy protocol at the Cleveland Clinic and sliced (1 cm thick) using a guided box. Slices were either rapidly frozen for biochemical analysis or short-fixed in 4% PFA followed by sectioning for morphological studies.

## Author contributions

BCS and JLW conceptualized and designed the study. BCS designed, performed, and analyzed *in vitro* experiments. BCS and RAT performed *in vivo* experiments and prepared murine samples for analysis. AM generated IPA analysis from data collected by JLW and BCS. MLH aided in analysis and presentation of IPA data. BCS performed and analyzed murine and human histology. RD provided human tissue samples and provided training and critical analysis of human tissue labeling and characterization. BCS and JLW wrote and edited the manuscript.

## Acknowledgements

We thank Misha Psenicka for his assistance in mouse colony management, husbandry, and genotyping and Dr. Benjamin Shaw for insightful discussion. We also thank Dr. Daniel M. Rotroff for the mentorship of Arshiya Mariam and the Cleveland Clinic Genomics Core for performing the RNA-sequencing on astrocytes. This work was supported by National Institute of Neurological Disorders and Stroke (NINDS) R01 NS119178 (awarded to Jessica L. Williams), R01 NS123532 (awarded to Ranjan Dutta), R35 NS097303 (awarded to Bruce D. Trapp), and National MS Society RFA-2203-39228 (awarded to Jessica L. Williams), and CSU CD-CAVS training grant NIH T32 HL150389 (awarded to Brandon C. Smith).

## Supplemental Figure Legends

**Supplemental Figure 1. PD-1 agonism and PD-1/PD-L1 antagonism primarily impacts LNCs cocultured with astrocytes.** Murine astrocytes and LNCs were harvested and co-cultured in the presence of media alone, 10 ng/ml IFNγ, 100 nM PD-1/PD-L1 inhibitor, and/or 1.0 μg/ml PD-1 agonist for 48 h. Caspase 3/7 activity was measured in (**A**) LNCs cultured alone, (**B**) astrocytes co-cultured with LNCs following LNC removal, (**C**) astrocytes cultured alone, (**D**) and in a LNC cultured plate following LNC removal to serve as a blank/background control. Caspase 3/7 activity was normalized to cell number. (**E**) Primary murine LNCs were stimulated with and without 10 ng/ml IFNγ for 24 h and RNA transcript levels of *Cd274* and *Pdcd1* were assessed. Data are representative of 2 independent experiments with 3-4 technical replicates each. All data represent the mean ± SEM. *P < 0.05 by one-way ANOVA.

**Supplemental Figure 2. Gating strategy to assess recombination efficiency in *Ifngr1*^fl/fl^ *Aldh1l1*-Cre^ERT2+^ mice.** TdTomato *Ifngr1*^fl/fl^ *Aldh1l1*-Cre^ERT2+^ mice were generated and treated with tamoxifen for 5 consecutive days. Spinal cord tissue was then digested and labeled with ACSA-2 to determine recombination efficiency. Cells were gated for singlets, live cells, ACSA-2 positivity, and then TdTomato positive cells.

**Supplemental Figure 3. EAE lesion characterization in *Ifngr1*^fl/fl^ *Aldh1l1*-Cre^ERT2+^ mice.** EAE was induced in *Ifngr1*^fl/fl^ *Aldh1l1*-Cre^ERT2+^ mice (*n* = 7) and *Ifngr1*^fl/fl^ littermate controls (*n* = 8) and EAE clinical course was blindly monitored. On day 16 ± 1 mice were injected i.p. with tamoxifen for 5 consecutive days to induce recombination. 35 days post-immunization, mice were perfused and the CNS was removed and cryopreserved for IHC analysis. Ventral white matter tracts of the lumbar spinal cord were imaged using confocal microscopy. (**A**) *Ifngr1*^fl/fl^ and (**B**) *Ifngr1*^fl/fl^ *Aldh1l1*-Cre^ERT2+^ tissue sections were labeled for MBP and nuclei were counterstained with DAPI. (**C**) Lesion area and (**D**) MBP positive area were quantified. Data represent the combined mean ± SEM from 2 independent experiments and were analyzed using a two-tailed Student’s *t* test. *P < 0.05, ***P < 0.001.

## Notes

### Competing Interest Statement

The authors have declared no competing interest.

