## Supplemental Figures for "Astrocyte interferon-gamma signaling dampens inflammation during chronic central nervous system autoimmunity via PD-L1"

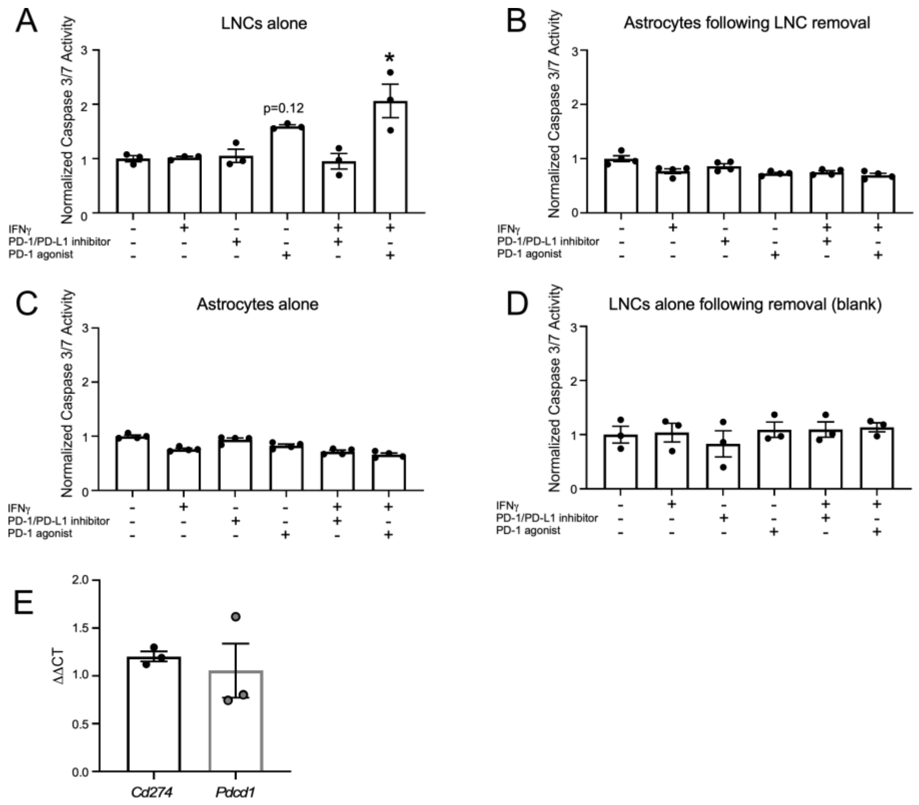


**Supplemental Figure 1. PD-1 agonism and PD-1/PD-L1 antagonism primarily impacts LNCs cocultured with astrocytes.** Murine astrocytes and LNCs were harvested and co-cultured in the presence of media alone, 10 ng/ml IFNγ, 100 nM PD-1/PD-L1 inhibitor, and/or 1.0 μg/ml PD-1 agonist for 48 h. Caspase 3/7 activity was measured in (**A**) LNCs cultured alone, (**B**) astrocytes co-cultured with LNCs following LNC removal, (**C**) astrocytes cultured alone, (**D**) and in a LNC cultured plate following LNC removal to serve as a blank/background control. Caspase 3/7 activity was normalized to cell number. (**E**) Primary murine LNCs were stimulated with and without 10 ng/ml IFNγ for 24 h and RNA transcript levels of *Cd274* and *Pdcd1* were assessed. Data are representative of 2 independent experiments with 3-4 technical replicates each. All data represent the mean ± SEM. *P < 0.05 by one-way ANOVA.

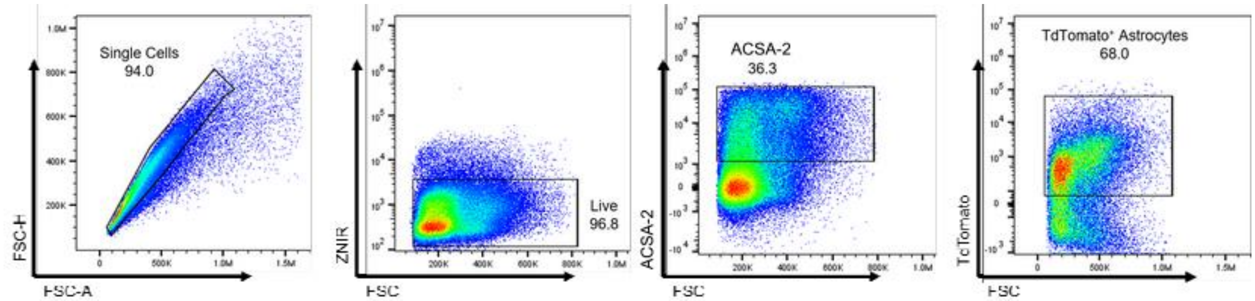


**Supplemental Figure 2. Gating strategy to assess recombination efficiency in *Ifngr1*$^{fl/fl}$ *Aldh1l1*-Cre$^{ERT2+}$ mice.** TdTomato *Ifngr1*$^{fl/fl}$ *Aldh1l1*-Cre$^{ERT2+}$ mice were generated and treated with tamoxifen for 5 consecutive days. Spinal cord tissue was then digested and labeled with ACSA-2 to determine recombination efficiency. Cells were gated for singlets, live cells, ACSA-2 positivity, and then TdTomato positive cells.

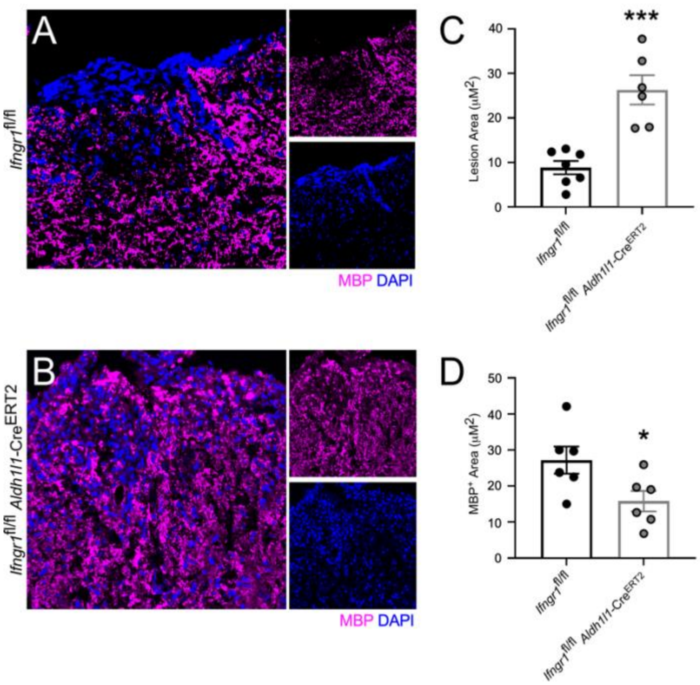


**Supplemental Figure 3. EAE lesion characterization in *Ifngr1*$^{fl/fl}$ *Aldh1l1*-Cre$^{ERT2+}$ mice.**

EAE was induced in *Ifngr1*$^{fl/fl}$ *Aldh1l1*-Cre$^{ERT2+}$ mice ($n = 7$) and *Ifngr1*$^{fl/fl}$ littermate controls ($n = 8$) and EAE clinical course was blindly monitored. On day 16 ± 1 mice were injected i.p. with tamoxifen for 5 consecutive days to induce recombination. 35 days post-immunization, mice were perfused and the CNS was removed and cryopreserved for IHC analysis. Ventral white matter tracts of the lumbar spinal cord were imaged using confocal microscopy. (**A**) *Ifngr1*$^{fl/fl}$ and (**B**) *Ifngr1*$^{fl/fl}$ *Aldh1l1*-Cre$^{ERT2+}$ tissue sections were labeled for MBP and nuclei were counterstained with DAPI. (**C**) Lesion area and (**D**) MBP positive area were quantified. Data represent the combined mean ± SEM from 2 independent experiments and were analyzed using a two-tailed Student's *t* test. $^{*}P < 0.05$, $^{***}P < 0.001$.
